# TrkB induced metastatic potential of cancer by suppression of BMP mediated tumor inhibitory activity

**DOI:** 10.1101/2020.01.29.924324

**Authors:** Min Soo Kim, Wook Jin

## Abstract

Our previous observations also demonstrate that TrkB expression in breast cancer induces metastatic potential by both JAK2/STAT3 and PI3K/AKT activation and induced metastasis of breast cancer mediated suppression of RUNX3 and KEAP1 expression by TrkB. Also, TrkB induced metastatic potential of cancer and suppressed the growth inhibitory activity in response to BMP signaling by preventing BMRRI/BMPRII complex formation. The previous report BMP-2 and BMP4 trigger tumor inhibitory activity in colorectal cancer by upregulation of RUNX3 expression. Although TrkB may regulate tumor inhibitory activity by BMP-induced upregulation of RUNX3, it is not still fully understood how TrkB signaling adjusts to inhibit BMP signaling-mediated tumor suppression.Our findings provide important molecular insights into TrkB-mediated modulation of BMP signaling has remained unknown, and none of the studies still reported a correlation between TrkB and BMP signaling. Our current study surprisingly showed that unique role of TrkB in the regulation of BMP-induced tumor inhibitory activity and BMP-2-induced RUNX3 expression.

## Introduction

The bone morphogenetic proteins (BMPs) family, which are consists of over 20 types of BMPs, are one of transforming growth factor-β (TGF-β) superfamily and are multifunctional growth factors with a broad range of regulatory functions, including embryonic development, apoptosis, chemotaxis, proliferation, and cellular differentiation [1-4]. As TGF-β signaling, secreted BMP-2 signals through a complex of BMP type II (BMPRII) and BMP type I (BMPRIA) receptor on the cell surface leads to activation of SMAD1 and thereby facilitates the nuclear translocation of SMAD1 in association with SMAD4 to regulate specific target gene expression [5-7].

Like functions of TGF-β in cancer microenvironments, BMPs and their receptors have implicated an essential role as a dual function, which has a tumor suppressor and promoter of tumor progression in cancer. For example, Gremlin 1 expression, BMP-antagonist, in an analysis of large patient cohorts significantly increased in tumor tissues of breast cancer patients [8]. Also, Gremlin 1 induces tumorigenesis of cancer cells by inhibiting BMP/SMAD signaling and associated poor prognosis of breast cancer patients [9]. Moreover, estrogen treatment increases the reduction of BMPRI [10]. Furthermore, expression of BMPRI or BMPRII exhibited significant loss in patients of poorly differentiated prostate cancer, renal cell carcinoma, urinary bladder cancer [5, 11, 12] and BMPRI expression frequently reduced and associated with poor survival in pancreatic cancer patients and inactivation of BMP signaling induces invasiveness of tumor [6] and BMP-2 induced apoptosis of glioma by induction of BMPRI [13]. However, other studies suggest that BMP signaling engaged in tumorigenesis and metastasis of tumors as tumor promotor. For instance, inhibition of BMP signaling by the antagonist of the BMPRIs (alk2, alk3, and alk6) decreases the growth and eventually promotes cell death in lung cancer cells [14]. Also, loss of BMPRIB associated with poor survival of breast cancer patients and render to the acquirement of drug-resistance [15]. So, it is still not completely clear what is a function of BMP signaling in tumor progression.

The recent studies, including our studies, demonstrated that BDNF and its TrkB receptor crucial role in tumorigenesis and metastasis of cancer and associated with poor survival of various cancer patients. BDNF/TrkB signaling activated in various cancer types, including breast cancer [16], colon cancer [17], lung cancer [18], pancreatic cancer [19], cutaneous melanoma [20], ovarian cancer [21], oral squamous cell carcinoma (OSCC). Also, Its expression associated with poor prognosis of melanoma and OSCC patients [22]. Moreover, TrkB inhibition induces suppression of survival of medulloblastoma by induction of apoptosis [23]. Our previous observations also demonstrate that TrkB expression in breast cancer induces metastatic potential by both JAK2/STAT3 and PI3K/AKT activation [24] and induced metastasis of breast cancer mediated suppression of RUNX3 and KEAP1 expression by TrkB [16]. The previous report described BMP signaling have linked to regulation of RUNX3 expression, a tumor suppressor. BMP-2 and BMP4 trigger tumor inhibitory activity in colorectal cancer by upregulation of RUNX3 expression [25]. Although TrkB may regulate tumor inhibitory activity by BMP-induced upregulation of RUNX3, it is not still fully understood how TrkB signaling adjusts to inhibit BMP signaling-mediated tumor suppression.

We show here that TrkB induced metastatic potential of cancer and suppressed the growth inhibitory activity in response to BMP signaling by preventing BMRRI/BMPRII complex formation.

## Results

### TrkB expression drives inhibition of established BMP signaling

To understand the correlation between TrkB expression and BMP signaling, we examined whether TrkB expression modulates BMP response element (BRE) luciferase activity. The transient transfection of TrkB in RIE-1, HeLa, and NMuMG cells inhibits BRE-luciferase activity relative to that of control (**Figures 1A, 1B**, and **S1A**). Thus, TrkB may inhibit BMP signaling for the acquisition of invasion and tumorigenesis of a tumor. To test this hypothesis, we generated RIE-1 and HeLa cells which overexpressed TrkB (Data not shown).

**Figure 1.**
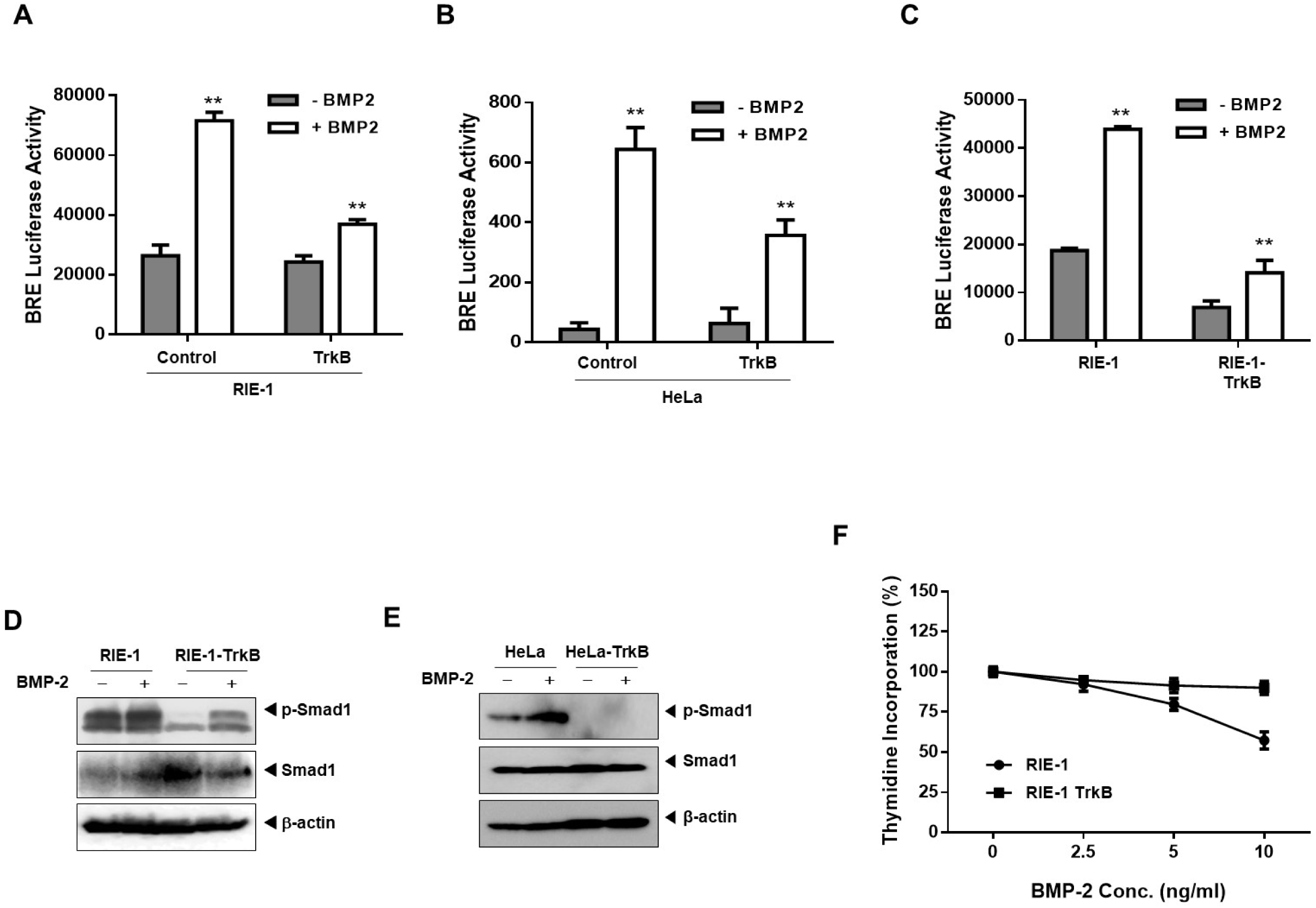
TrkB overexpression induces inhibition of BMP-2-mediated tumor suppressor activity. (A, B) The activity of BMP-2-responsive BRE Luciferase reporter in RIE-1 (A) and HeLa cells (B) transfected with the TrkB. Luciferase activity was measured 24 h after treatment with BMP-2 (5 ng/mL). **Control versus treatment with BMP-2, *P* < 0.05. n = 3. (C) The activity of BMP-2-responsive BRE Luciferase reporter in RIE-1 and RIE-1-TrkB cells. **Control versus treatment with BMP-2, *P* < 0.05. n = 3. (D) Western blot analysis of the expression of phospho-SMAD1 and SMAD1 in RIE-1, HeLa, or RIE-1-TrkB, HeLa-TrkB cells after stimulation with BMP-2 (5 ng/mL). (E) Thymidine incorporation assay of RIE-1 or RIE-1-TrkB cells treated with various concentrations of BMP-2 as indicated. Points, averages of means from three determinations; bars, SD.

As expected, BMP-2-induced transcriptional activity in RIE-1-TrkB and NMuMG-TrkB cells reduced by TrkB overexpression (**Figures 1C** and **S1B**). In normal RIE-1 and HeLa cells, BMP-2-induced phosphorylation of SMAD1 was readily detectable but increased SMAD1 phosphorylation markedly reduced in RIE-1- and HeLa-TrkB cells relative to RIE-1 and HeLa cells (**Figures 1D** and **1E**). To further understand the function of TrkB on the BMP-2 signaling, we then investigated whether TrkB suppresses BMP-2-induced growth inhibition. RIE-1-TrkB cells were resistant to BMP-2-induced growth inhibition. RIE-1 cells failed to resistant to BMP-2-induced growth inhibition observed in RIE-1-TrkB cells (**Figure 1F**). These results suggest that TrkB expression inhibits BMP-2-induced growth inhibitory activity.

### TrkB promotes the ability of tumorigenicity and Metastasis of RIE-1 Cells

The above results suggest that TrkB expression may enhance the tumorigenic potential of the tumor by BMP-2-mediated growth inhibition. Based on the above findings, we evaluated whether TrkB overexpression leads to the induction of metastatic potential to RIE-1 cells. To metastasize to distant organs, cancer cells have to overcome anoikis during cancer progression [26]. We first examined whether TrkB confers anoikis resistance to achieve invasion and dissemination, which is the hallmark for metastatic cancer. RIE-1-TrkB cells form large aggregates in suspension and promote cell anchorage-independent growth relative to RIE-1 cells (**Figure 2A**). In cell migration assay, RIE-1-TrkB cells had markedly increased cell migration, but RIE-1 cells significantly decreased cell migration, indicating that the increased cell migration correlated with the level of TrkB expression (**Figure 2B**). We then investigated whether TrkB expression affected the colony-forming ability of RIE-1 cells. RIE-1-TrkB cells increase 3 ∼ 4 fold higher colony-forming activity relative to RIE-1 cells (**Figure 2C**). Next, we performed an in vitro mammosphere forming assay, which is associated with mammary stem-like traits [27]. Relative to the parental RIE-1 cells, RIE-1-TrkB cells were 2.6 fold enriched in mammosphere-forming cells (**Figure 2D**). Also, in wound healing assay, TrkB overexpression in RIE-1 cells significantly increases cell motility compared with that in parental RIE-1 cells (**Figure 2E**), indicating that TrkB expression enhances tumor outgrowth. Also, these findings suggest that TrkB accelerates the invasion-metastasis cascade for disseminating to distant organ sites.

**Figure 2.**
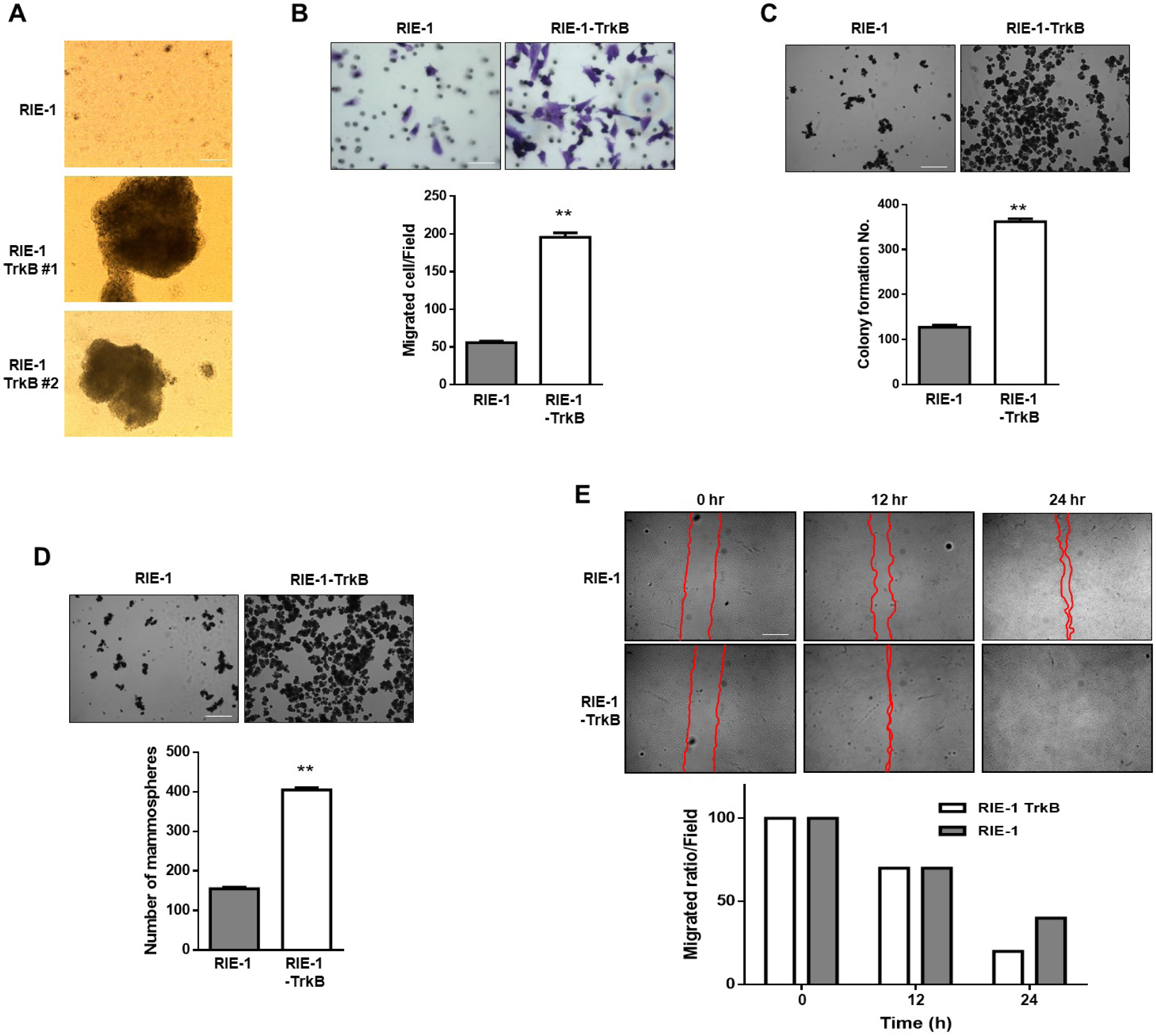
TrkB induces cell migration, survival, and the transforming activity of tumor cells. (A) Formation of spheroid colonies of RIE-1 or RIE-1-TrkB cells. Spheroid colonies in the suspension then photographed at ×200 magnification. (B) The migration assay: bright-phase microscopy images and quantification of RIE-1 or RIE-1-TrkB cells (n = 3). P < 0.0001, t-test. (C) Colony-forming assay: bright-phase microscopy images and quantification of RIE-1 or RIE-1-TrkB cells (n = 3). P < 0.0001, t-test. (D) Mammosphere assay: bright-phase microscopy images and quantification of RIE-1 or RIE-1-TrkB cells (n = 3). P < 0.0001, t-test. (E) Wound-healing assay: bright-phase microscopy images and quantification of RIE-1 or RIE-1-TrkB cells.

### Loss of TrkB restore BMP-mediated tumor inhibitory activity

To further substantiate the regulation of BMP signaling in the tumor invasion function of TrkB, we also knocked down the expression of TrkB in MDA-MB-231 and Hs578T cells and found that knockdown of TrkB led to a drastically increases BMP-2-induced BRE transcriptional activity relative to control-shRNA cells (**Figures 3A** and **3B**). We also observed SMAD1 phosphorylation in MDA-MD-231 TrkB-shRNA cells being increased after BMP-2 stimulation, whereas the treatment of BMP-2 fails to induction of phosphorylation of SMAD1 in MDA-MB-231 control-shRNA cells (**Figure 3C**). We then examined whether TrkB knockdown restored BMP-2-induced growth inhibition. MDA-MB-231 control-shRNA cells were resistant to BMP-2-induced growth inhibition, whereas this resistance to BMP-2-mediated growth inhibition severely compromised in TrkB knockdown cells (**Figure 3D**). These results indicate that TrkB plays a role in suppressing BMP-2-mediated growth inhibitory activity for supporting tumor invasion and metastasis.

**Figure 3.**
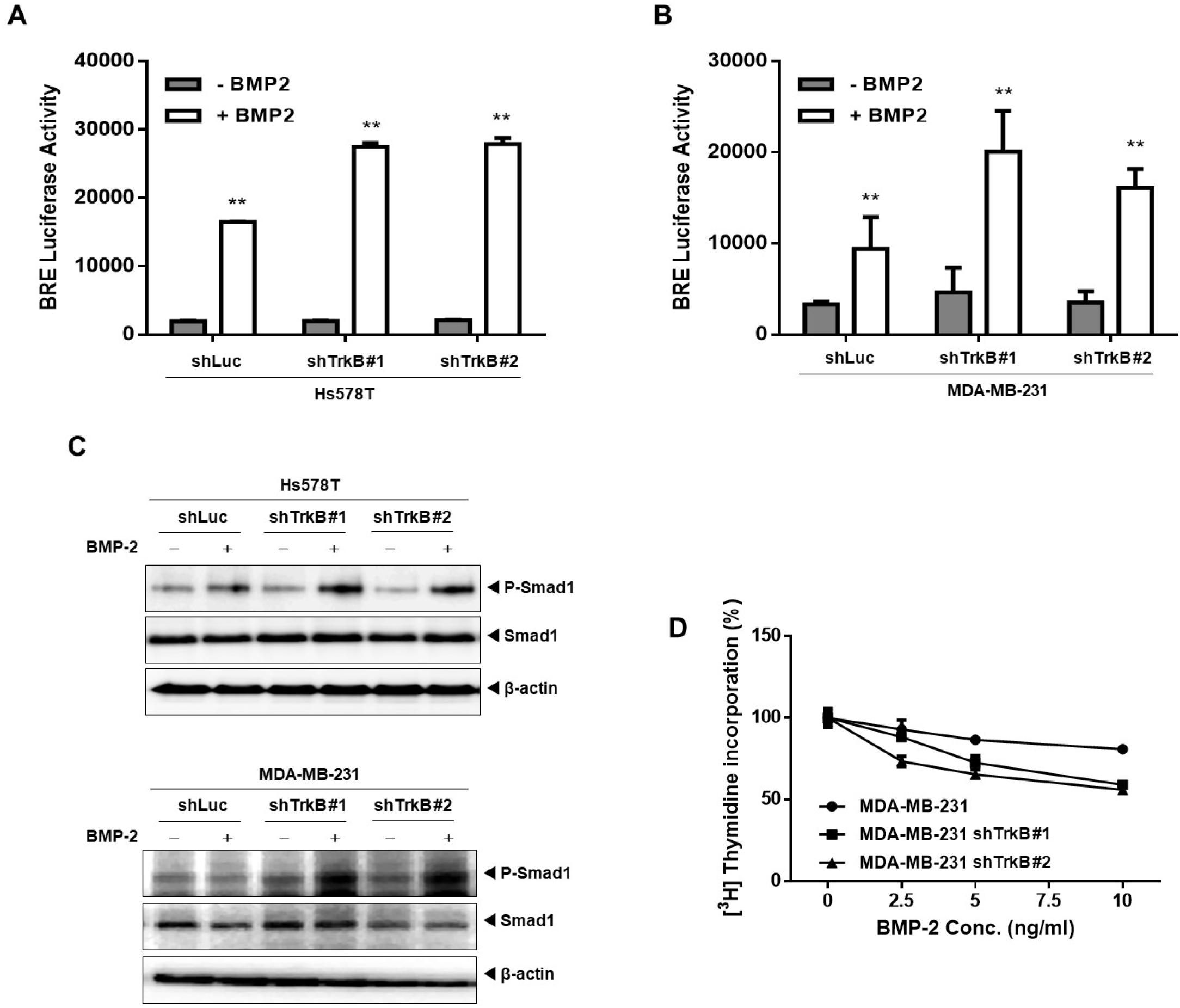
loss of TrkB in highly metastatic breast cancer cells restore inhibited BMP signaling. (A) The activity of BMP-2-responsive BRE Luciferase reporter in Hs578T control-shRNA or TrkB-shRNA cells. Luciferase activity was measured 24 h after treatment with BMP-2 (5 ng/mL). **Control versus treatment with BMP-2, *P* < 0.05. n = 3. (B) The activity of BMP-2-responsive BRE Luciferase reporter in MDA-MB-231 control-shRNA or TrkB-shRNA cells. **Control versus treatment with BMP-2, *P* < 0.05. n = 3. (C) Western blot analysis of the expression of phospho-SMAD1 and SMAD1 in Hs578T and MDA-MB-231 control-shRNA or TrkB-shRNA cells after stimulation with BMP-2 (5 ng/mL). (D) Thymidine incorporation assay of MDA-MB-231 control-shRNA or TrkB-shRNA cells treated with various concentrations of BMP-2 as indicated. Points, averages of means from three determinations; bars, SD.

### TrkB directly interacts with BMP type II receptor to inhibits BMP signaling

Our above results suggest that TrkB may suppress BMP-mediated tumor inhibitory activity through the regulation of upstream BMP receptors. To unravel the mechanism of TrkB-mediated BMP-2 signaling regulation, we speculated that TrkB might suppress BMP signaling via interacting with BMP receptors. So, we sought to identify proteins that distinctly recognized by TrkB in BMP-2-mediated tumor inhibitory activity. We found TrkB directly interacts with BMPRII, whereas TrkB does not interact with BMPRI (**Figures 4A** and **4B**). We then examined the endogenous interaction between TrkB and BMPRII in MDA-MB-231 cells, which express TrkB and tissues of breast cancer patients. Endogenous TrkB in MDA-MB-231 cells directly interacted with BMPRII and upregulation of TrkB exhibited in tumor tissue of the breast cancer patients relative to healthy tissue and interaction between endogenous TrkB, and BMPRII also confirmed by immunoprecipitation (**Figures 4C** and **4D**). Moreover, The deletion of the cytoplasmic domain and N terminal region, including the extracellular and transmembrane domain of BMPRII did not change on this binding, but the elimination of the area between 203 amino acid and 350 amino acid in the tyrosine kinase domain of BMPRII abolished this binding (**Figure 4E**). Thus, the kinase domain of BMPRII mediates its interaction with TrkB.

**Figure 4.**
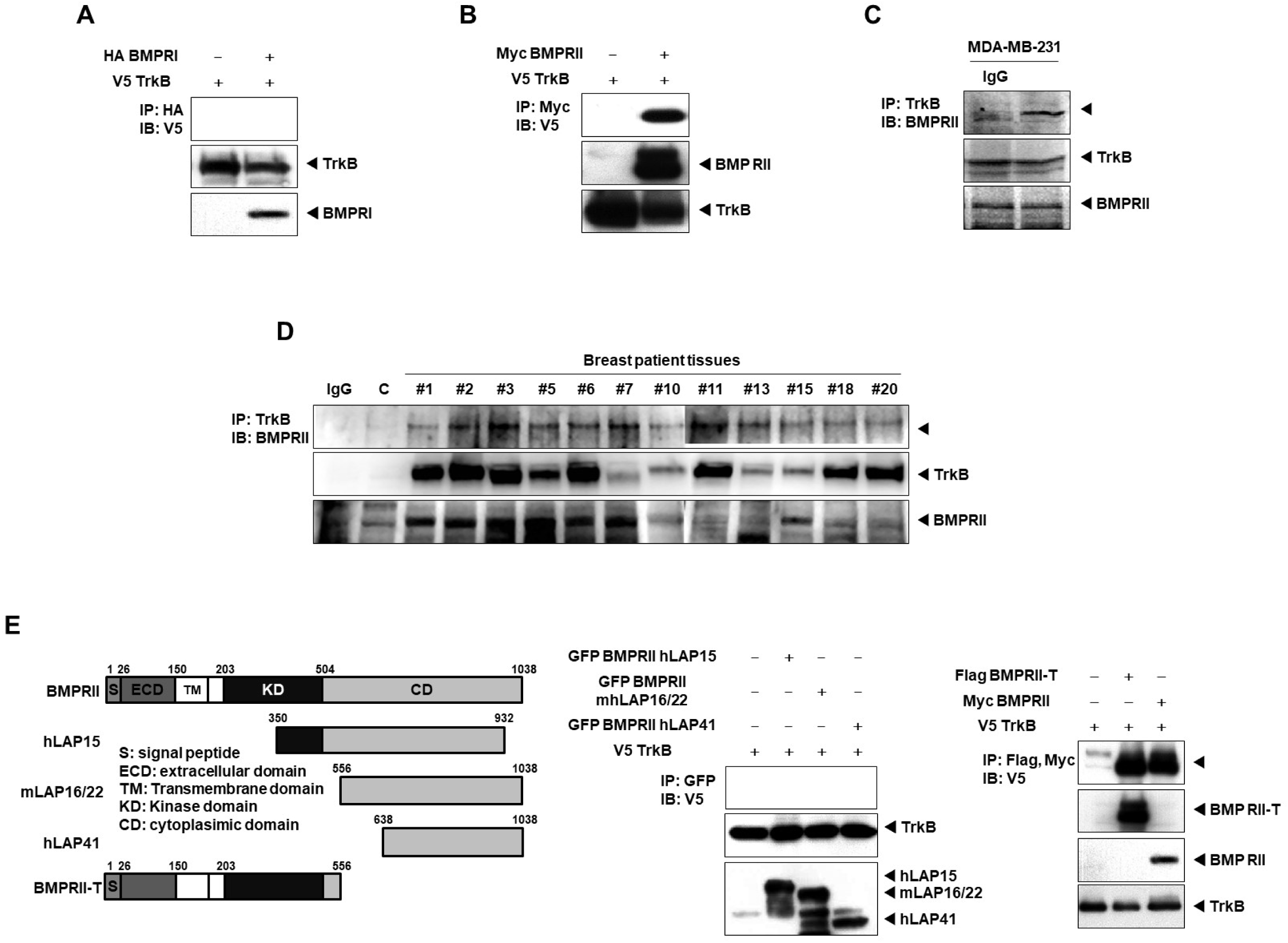
TrkB inhibits BMP signaling via block interaction between BMPRI and BMPRII in response to BMP-2. (A) Western blot analysis of the whole-cell lysates and immunoprecipitates of HA derived from 293T cells transfected with the V5-TrkB and HA-BMPRI constructs, as indicated. (B) Western blot analysis of the whole-cell lysates and immunoprecipitates of Myc derived from 293T cells transfected with the V5-TrkB and Myc-BMPRII constructs, as indicated. (C) Identification of the complex formation of endogenous TrkB-BMPRII in MDA-MB-231 cells. Western blot analysis of the whole-cell lysates and immunoprecipitates of TrkB derived from MDA-MB-231 cells, as indicated. (D) Identification of the complex formation of endogenous TrkB-BMPRII in tumor tissues of breast cancer patients. Western blot analysis of the whole-cell lysates and immunoprecipitates of TrkB derived from tumor tissues of breast cancer patients, as indicated. (E) Identification of the BMPRII region that interacted with TrkB. Western blot analysis of whole-cell lysates and immunoprecipitates of GFP, Flag, or Myc derived from 293T cells transfected with V5-TrkB, Myc-BMPRII, and the BMPRII deletion construct as indicated.

### The tyrosine kinase activity of TrkB required for inhibition of BMP-2 signaling

We assessed the effects of TrkB kinase inhibitor in BMP-mediated tumor inhibitory activity in RIE-1 and RIE-1-TrkB cells. Inhibiting TrkB kinase activity with K252a significantly increases BMP-2-induced BRE transcriptional activity in RIE-1-TrkB cells, whereas there was no change induction of BMP-2-mediated BRE luciferase activity in RIE-1 cell with or without treatment of K252a (**Figure 5A**). Also, similar results obtained with the thymidine incorporation assay. The ability of TrkB to resistant in response to BMP was significantly increased in RIE-1-TrkB cells, whereas RIE-1-TrkB cells with treatment of K252a failed to rescue the ability of TrkB resistant to the growth inhibitory property of BMP (**Figure 5B**).

Similarly, MDA-MB-231 and Hs578T breast cancer cells in which expression of the endogenous TrkB more increase reporter activity in response to K252a and BMP-2 (**Figures 5C** and **5D**). Consistent with rescued BMP-induced BRE-luciferase activity and growth inhibitory effect, the level of SMAD1 phosphorylation in RIE-1-TrkB cells by treatment of K252a significantly increased relative to control with treated k252a in response to BMP-2 (**Figure 5E**). Also, we were intrigued about the K252a effect to inhibition of BMP-2 signaling in RIE-1-TrkB cells because it implies that activation of TrkB by overexpression may be required for suppression of BMP signaling. We, therefore, validated the possibility that TrkB is activated by overexpression via a coimmunoprecipitation experiment. Our results exhibited that RIE-1-TrkB cells indeed was dramatically increased the phosphorylation level of TrkB regardless with or without BDNF treatment relative to RIE-1 cells (**Figure S2**). These results exhibit that TrkB takes action in inhibition of BMP-induced growth inhibitory effect after activation of TrkB.

**Figure 5.**
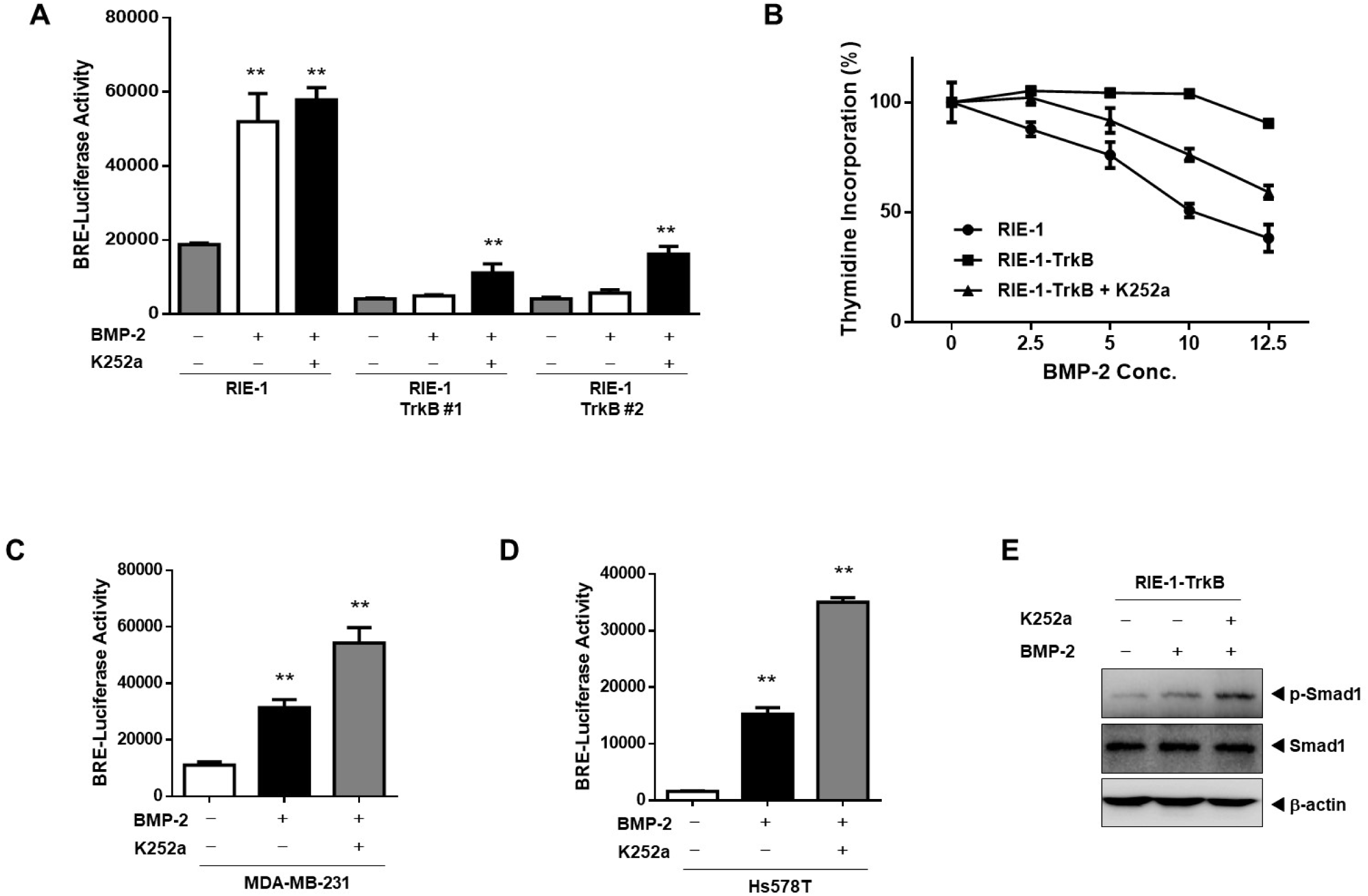
The tyrosine kinase activity of TrkB requires inhibiting BMP signaling. (A) The activity of BMP-2-responsive BRE Luciferase reporter in RIE-1 and RIE-1-TrkB cells in response to K252a (50nM). Luciferase activity was measured 24 h after treatment with BMP-2 (5 ng/mL). **Control versus treatment with BMP-2, *P* < 0.05. n = 3. (B) Thymidine incorporation assay of RIE-1 and RIE-1-TrkB cells treated K252a (50nM) with various concentrations of BMP-2 as indicated. Points, averages of means from three determinations; bars, SD. (C, D) The activity of BMP-2-responsive BRE Luciferase reporter in MDA-MB-231 and Hs578T cells in response to K252a (50nM). **Control versus treatment with BMP-2, *P* < 0.05. n = 3. (E) Western blot analysis of the expression of phospho-SMAD1 and SMAD1 in RIE-1 and RIE-1-TrkB cells with or without BMP-2 (5 ng/mL) or K252a (50nM).

To more assess the effects of activation of TrkB in BMP-induced transcriptional activity, we generated RIE-1 TrkB KD cells using vectors that expressing kinase-dead mutant of TrkB (K588M). As expected, the BRE-luciferase activity of RIE-1-TrkB or HeLa-TrkB cells drastically inhibits reporter activity, but the expression of TrkB KD in RIE-1 and HeLa cells fail to suppress reporter activity in response to BMP-2 (**Figures 6A** and **6B**). Also, although TrkB expression reduced the phosphorylation level of SMAD1, the TrkB K588M failed to do so (**Figure 6C**). These results also suggest that the activation of TrkB by overexpression is sufficient for supporting tumor invasion without BDNF through inhibiting BMP-mediated tumor inhibitory activity.

**Figure 6.**
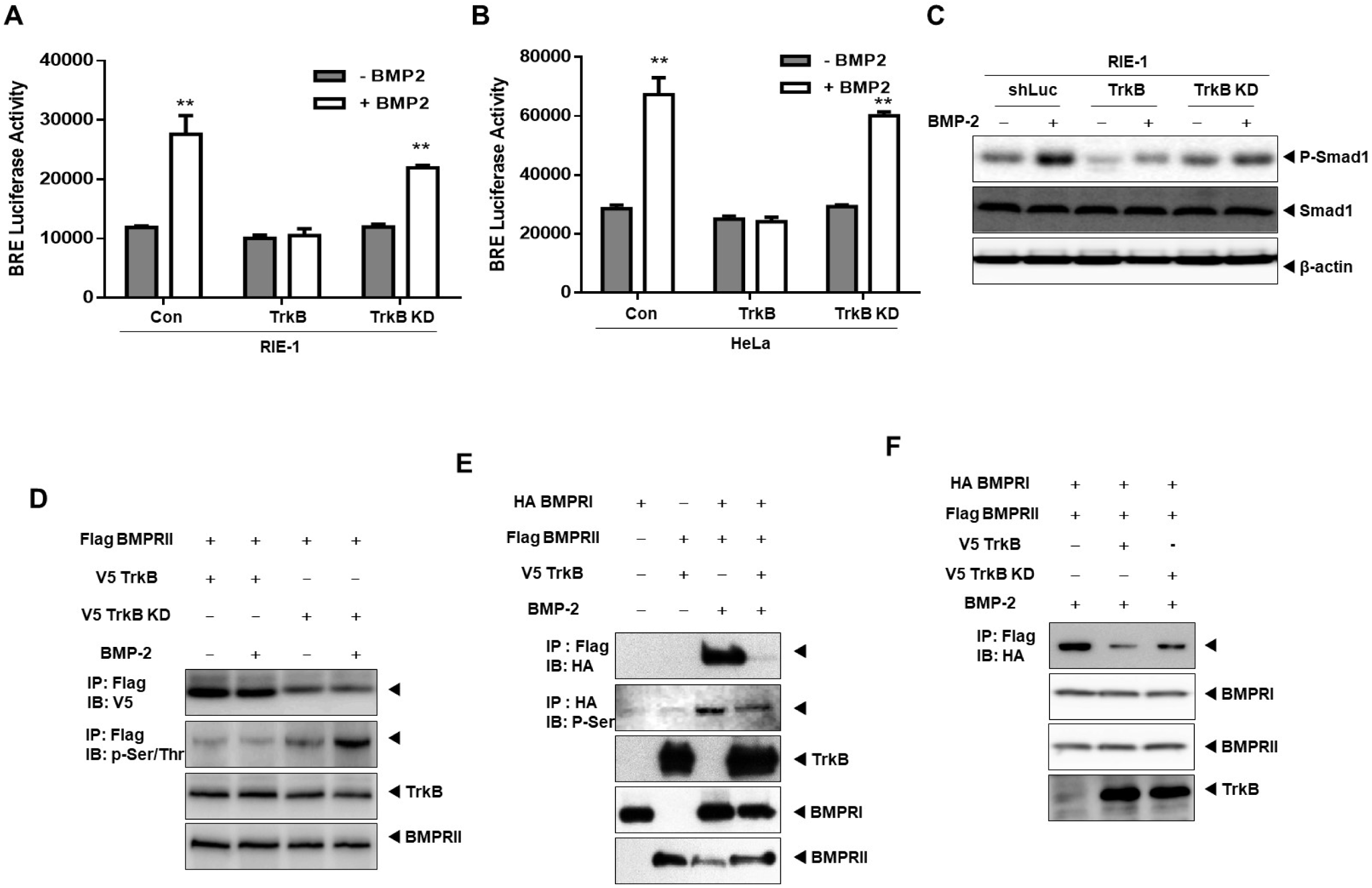
The loss of kinase activity of TrkB restores BMP signaling by enhancing interaction between BMPRI and BMPRII in response to BMP-2. (A, B) The activity of BMP-2-responsive BRE Luciferase reporter in RIE-1 and HeLa cells transfected with the TrkB, or RIE-1-TrkB KD constructs. **Control versus treatment with BMP-2, *P* < 0.05. n = 3. (C) Western blot analysis of the expression of phospho-SMAD1 and SMAD1 in RIE-1, RIE-1-TrkB, and RIE-1-TrkB KD cells with or without BMP-2 (5 ng/mL). (D) Western blot analysis of whole-cell lysates and immunoprecipitates of Flag derived from 293T cells after transfection of V5-TrkB, V5-TrkB KD, or Flag-BMPRII constructs with or without BMP-2 treatment. (E) Western blot analysis of whole-cell lysates and immunoprecipitates of Flag derived from 293T cells after transfection of HA-BMPRI, Flag-BMPRII, or V5-TrkB constructs with or without BMP-2 treatment. (F) Western blot analysis of whole-cell lysates and immunoprecipitates of Flag derived from 293T cells after transfection of HA-BMPRI, Flag-BMPRII, V5-TrkB, or V5-TrkB KD constructs with BMP-2 treatment.

Our attention also is drawn to possible TrkB KD-BMPRII complex formation. TrkB wild type strongly associated with BMPRII, but the interaction between BMPRII and TrkB KD mutant significantly decreased with or without BMP-2 treatment, and the phosphorylation of BMPRII by TrkB KD mutant restored in response to BMP-2 relative to TrkB wild type (**Figure 6D**). Because the interaction between BMPRI and BMPRII enhanced by stimulation of BMP-2, we speculated that BMPRII-TrkB complex formation interferes interaction between BMPRI and BMPRII in the presence of BMP-2. The result showed that BMPRII could strongly bind to BMPRI by treatment of BMP-2, but this binding completely abolished in the presence of TrkB. Also, the increased phosphorylation level of BMPRI after treated BMP-2 markedly decreased by TrkB (**Figure 6E**). Moreover, the TrkB-mediated depletion of BMPRII-BMPRI interaction rescued by TrkB kinase-dead mutant (**Figure 6F**), indicating that activation of TrkB required for tumorigenesis via inhibition of BMP signaling.

### TrkB regulates expression of BMP type I receptor to inhibits BMP signaling

BMP-2 induces expression of RUNX3 as a tumor suppressor and subsequently decreases c-Myc expression via binding induced RUNX3 to the promotor of c-Myc. Also, dominant-negative BMPRI (DN-ALK3) in the presence of BMPRII does not suppress c-Myc transcriptional activity relative to the transfection of constitutively-active BMP Receptor IA (CA-ALK3) [25], indicating that activation of BMP signaling by phosphorylation or expression of BMPRI may be required for the RUNX3-induced inhibition of c-Myc expression. Previously, we showed that TrkB promotes tumorigenesis and metastasis of breast cancer through suppression of RUNX3 expression [28], and our present result showed that TrkB does not interact with BMPRI (**Figure 4A**). However, it is still not completely clear how TrkB adjusts to suppress RUNX3 expression by inhibiting BMP signaling. We speculate that TrkB may inhibit BMP-induced tumor growth inhibitory activity through the depletion of RUNX3 expression via regulation of BMPRI expression. To unravel to identify the mechanism of TrkB-mediated suppression of RUNX3 expression in BMP signaling, we examined the expression of BMPRI and BMPRII. The level of mRNA and protein expression of BMPRI severely compromised in Hela-TrkB and RIE-1-TrkB cells relative to Hela and RIE-1 cells (**Figures 7A, S3A**, and **S3B**), but TrkB does not affect BMPRII expression (**Figure S3B**). Inhibiting TrkB kinase activity by mutation of the kinase domain of TrkB effectively rescued loss of BMPRI expression by TrkB but does not affect BMPRII expression in both RIE-1-TrkB and RIE-1-TrkB KD cells (**Figures 7B** and **S3D**). Also, knockdown of TrkB in MDA-MB-231 and Hs578T cells induced BMPRI expression (**Figures 7C, 7D, 7E, 7F**, and **S3C**) but not BMPRII (Figures **S3E** and **S3G**). Moreover, we examined the level of BMPRI expression in the lungs of mice carrying MDA-MB-231 control-shRNA or TrkB-shRNA cells. BMPRI expression in the lungs of mice with MDA-MB-231 control-shRNA cells was significantly lower than that of the MDA-MB-231 TrkB-shRNA cells (**Figure 7G**). However, BMPRII expression showed no changed (**Figure S3G**).

**Figure 7.**
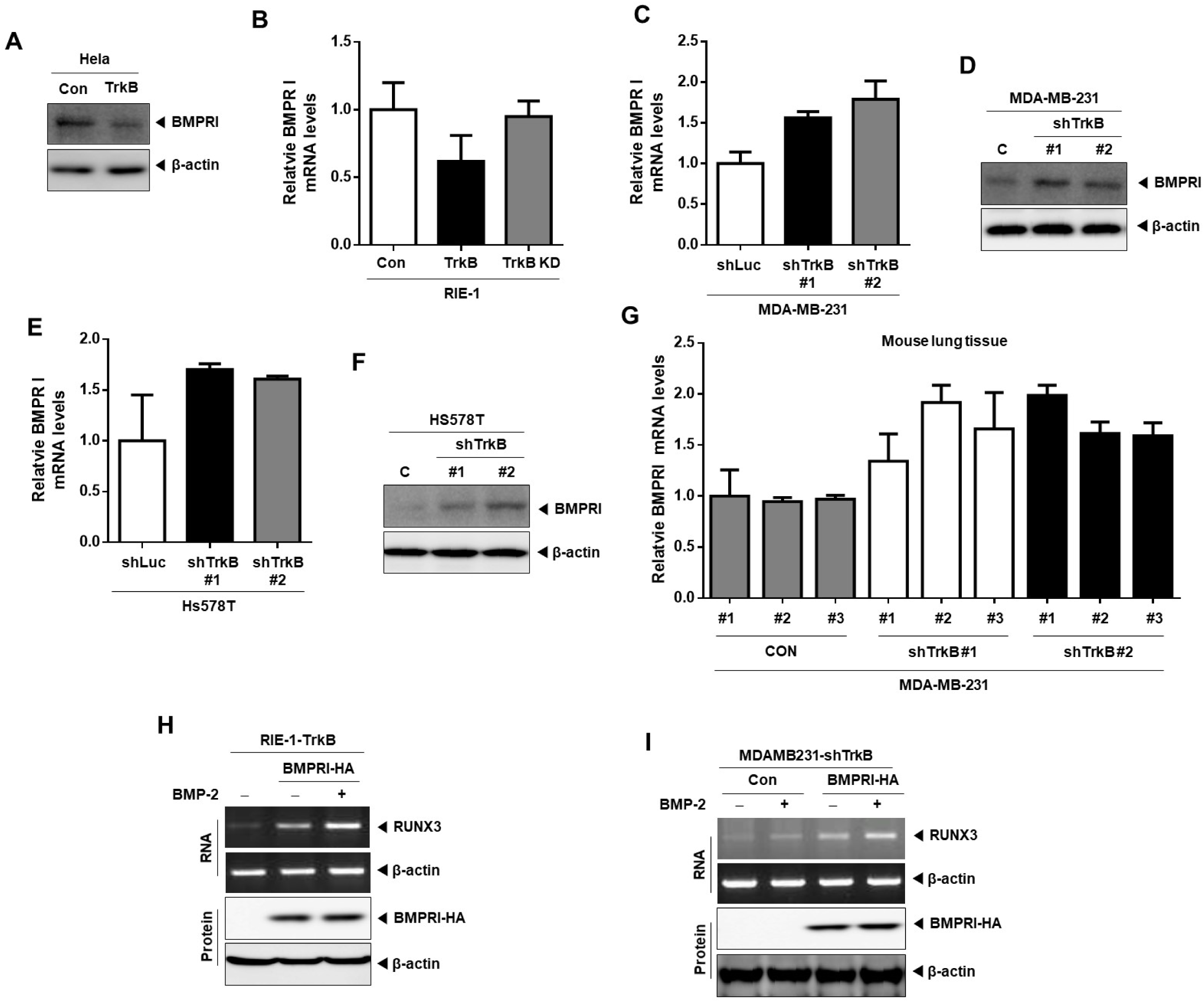
TrkB suppresses BMP-induced RUNX3 expression through the depletion of BMPRI expression. (A) Western blot analysis of the expression of BMPRI in HeLa and HeLa-TrkB cells. Loading control, β-actin. (B) The relative expression levels of BMPRI mRNA in RIE-1, RIE-1-TrkB, and RIE-TrkB KD cells, as determined by quantitative RT-PCR. Loading control, 18S. P < 0.05, t-test. (C) The relative expression levels of BMPRI mRNA in MDA-MB-231 control-shRNA and TrkB-shRNA cells, as determined by quantitative RT-PCR. Loading control, 18S. P < 0.05, t-test. (D) Western blot analysis of the expression of BMPRI in MDA-MB-231 control-shRNA and TrkB-shRNA cells. Loading control, β-actin. (E) The relative expression levels of BMPRI mRNA in Hs578T control-shRNA and TrkB-shRNA cells, as determined by quantitative RT-PCR. Loading control, 18S. P < 0.05, t-test. (F) Western blot analysis of the expression of BMPRI in Hs578T control-shRNA and TrkB-shRNA cells. Loading control, β-actin. (G) The relative expression levels of BMPRI mRNA in the lungs of individual mice harboring either MDA-MB-231 control-shRNA or TrkB-shRNA cells, as determined by quantitative RT-PCR. Loading control, 18S. P < 0.05, t-test. (H) The relative expression levels of RUNX3 mRNA and protein in RIE-TrkB cells transfected with BMPRI and treated with BMP-2 (I) The relative expression levels of RUNX3 mRNA and protein in MDA-MB-231 TrkB-shRNA cells transfected with BMPRI and treated with BMP-2

Our above result showed that complex formation between BMPRI and BMPRII significantly suppressed in the presence of TrkB (**Figures 6E** and **6F**). Under this condition, we observed whether BMPRI induce RUNX3 expression in response to BMP-2. Markedly upregulated RUNX3 expression observed by the introduction of BMPRI in RIE-TrkB cells and more induced by BMP-2 relative to RIE-1-TrkB cells (**Figure 6H**), but RUNX3 expression does not induce by transfection of BMPRII (**Figure S4A**). Also, RUNX3 expression in MAD-MB-231-TrkB shRNA cells more induced in response to BMP-2. Moreover, drastically upregulated RUNX3 expression observed by the introduction of BMPRI in MDA-MB-231 TrkB-shRNA cells and more induced by BMP-2 relative to MDA-MB-231 TrkB-shRNA cells (**Figure 6I**), but RUNX3 expression does not change by transfection of BMPRII (**Figure S4B**).

These results suggest that TrkB inhibits expression of RUNX3 by depleting BMPRI expression.

## Discussion

We find that TrkB that induce tumorigenesis and metastasis of cancer via induction of the JAK2/STAT3 pathway and PI3K/AKT pathway also governs the conversion to the suppression of BMP-mediated tumor inhibition. Our and another recent study demonstrated that TrkB expression drastically increased in triple-negative breast cancer [16] relative to other subtypes, and BDNF/TrkB plays a critical role in the generation and renewal of CSCs and recurrence of breast cancer after chemotherapy [29]. BDNF expression markedly increased in recurrent TNBCs isolated from patients after chemotherapy and promotes CSC self-renewal by increasing the ALDH1 expression. Moreover, TrkB+ CSCs exclusively overlapped with the ALDH1+ population, and treatment of BDNF in TrkB+/ALDH1+ cells induces KLF4 expression, which is one of stem cell markers. Furthermore, eradicating the TrkB+ CSCs increases survival by preventing of recurrence of TNBCs. These results suggest that TrkB is a critical regulator in the progression and recurrence of cancer.

BMP signaling prevents TGF-β-mediated CSCs transition in breast cancer and TGF-β-induced EMT in renal fibrosis. Chordin-like 2 and Gremlin 1, BMP antagonists, significantly increased, and BMP ligands such as BMP-2 and BMP4 markedly decreased in a mesenchymal subpopulation of human breast mammary epithelial cells. Also, BMP4 treatment markedly decreased cell migration and ability of mammosphere formation in a spontaneously arising mesenchymal subpopulation (MSP) of cells isolated from immortalized human MECs (HMLE), and Twist overexpressed HMLE cells [30, 31]. Moreover, Gremlin 1 promotes its stem cell maintenance in glioma and colorectal cancer [32, 33], and upregulation of Gremlin 1 significantly correlated with induced stem cell markers and poor survival of estrogen receptor (ER)-negative breast cancer patients [8]. These previous results are conspicuously consistent with our present results. Our current study surprisingly showed that TrkB has an unappreciated function in the inhibition of BMP signaling. BMP-2 induced SMAD1 phosphorylation and BMP-2-induced transcriptional activity in RIE-1 and HeLa cells significantly reduced by TrkB. In contrast, Loss of TrkB in highly metastatic breast cancer cells (MDA-MB-231 and Hs578T) restore BMP signaling. Moreover, BMP-2-induced growth inhibition significantly rescued by knockdown of TrkB in MDA-MB-231 and Hs578T cells. Our data suggest that TrkB also may inhibit canonical BMP signaling-mediated transcriptional activity.

Runt-related transcription factor 3 (RUNX3), a tumor suppressor, reduced in various cancer by mislocalization, hypermethylation, or loss of heterozygosity [34-36]. RUNX3 suppresses the function of nuclear-translocated Yes-associated protein (YAP) in cancer. Nuclear translocation of YAP (activation form) by dysregulation of the Hippo pathway has implicated in cancer progression and poor survival of patients [37-39]. RUNX3 inhibits YAP-induced EMT, tumorigenic, and metastatic potential of breast cancer by YAP-RUNX3 complex formation [40]. BMP signaling may involve inhibition of tumor development and progression by inducing the Hippo signaling pathway through upregulation of phospho-YAP (inactivation form) and stimulation of ras association domain family (RASSF1) as an upstream regulator of Hippo signaling pathway. BMP-2 treatment induces kinase cascade of Hippo signaling pathway, including upregulation of mammalian Ste20-like kinase 1 (MST1), MOB kinase activator 1 (MOB1), phospho-YAP expression, which behaves as a tumor suppressor. The upregulation of BMP-2 also induces BMPRII expression and promotes apoptosis by and inhibition of nuclear translocation of YAP and induction of RASSF1-MST1 complex formation via disruption of AKT-RASSF1 interaction [41]. Another report showed that canonical BMP signaling induces expression of RUNX3 and BMP2/4-induced RUNX3 suppress tumor growth through repression of c-Myc promoter activity via binging RUNX-binding elements in c-Myc promoter [25]. The potential linkage between BMP signaling and RUNX3 in these previous observations now can be accounted for by the mechanism presented in our results. Our previous and present studies surprisingly showed that TrkB represses RUNX3 expression by activation of the PI3K/AKT signaling pathway and inhibition of canonical BMP signaling. The introduction of BMPRI in the presence of TrkB upregulate RUNX3 expression, and its upregulation more increased by BMP-2. Also, RUNX3 expression increased in MDA-MB-231 TrkB-shRNA cells transfected with BMPRI relative to MDA-MB-231 control cells. Moreover, RUNX3 expression in MDA-MB-231 TrkB-shRNA cells transfected with BMPRI markedly increased by BMP-2.

So far, TrkB-mediated modulation of BMP signaling has remained unknown, and none of the studies still reported a correlation between TrkB and BMP signaling. Our current study surprisingly showed that unique role of TrkB in the regulation of BMP-induced tumor inhibitory activity and BMP-2-induced RUNX3 expression.

## Material and Methods

### Cell Lines, culture conditions and chemical inhibitors

RIE-1, HeLa, NMuMG, 293T, and human highly metastatic cancer cells (Hs578T and MDA-MB-231) cells were cultured in Dulbecco’s modified Eagle’s medium (DMEM) supplemented with 10 % fetal bovine serum (FBS). BMP-2 from PeproTech (500-P195BT) used at the final concentration of 50 ng/ml for the indicated time. The protein kinase inhibitor K252a purchased from Abcam.

### Plasmids and viral production

RIE-1, HeLa, and NMuMG cells were infected with a V5-TrkB using pLenti6.3/V5-TOPO TA Cloning Kit (Invitrogen) to generate TrkB overexpression cells [24] and and the TrkB K588M mutant using previously described [42] was generated by site-directed mutagenesis with Site-Directed Mutagenesis Kit (ThermoFisher Scientific). To generate a stable knockdown of TrkB, small hairpin-expressing vectors were purchased from Sigma-Aldrich (SHCLNG-NM_006180). Production and infection of target cells were previously described [24] and selected with 2 μg/ml puromycin and 500 μg/ml G418. Plasmid transfections were carried out using Lipofectamine 2000 (Invitrogen) reagent according to the manufacturer’s instructions.

### Human breast tumor samples

Proteins extracted from human breast normal and tumor samples obtained as previously described [16, 43].

### Antibodies, Western blotting, immunoprecipitation, and immunofluorescence

We performed Western blotting, immunoprecipitation, and immunofluorescence analysis, as previously described [44]. phospho-SMAD1 (ab214423), SMAD1 (ab63356), BMPRI, BMPRI, phosphor-tyrosine, phosphor-serine, and TrkB were from Abcam. Flag and β-actin were from Sigma-Aldrich; V5 was from Invitrogen; HA, Myc, and GFP were from Santa Cruz Biotechnology.

### Invasion, anoikis, wound healing, anchorage-independent cell growth, and mammosphere assays

All the assays performed as previously described [24, 45, 46]. For anchorage-independent cell growth and soft agar assays, 1 × 10^3^ cells/well seeded into 6 well cell culture plates. For the wound healing assay, 1 × 10^6^ cells/well seeded into 6 well cell culture plates. For the invasion assay, 1 × 10^4^ cells seeded into 24-well BD Matrigel invasion chambers with 8 μm pores (Corning, 62405-744). For anoikis assay, RIE-1 and RIE-1-TrkB cells were seeded into an Ultra Low Cluster plate (Corning) at 1 × 10 ^5^ cells per well in a six-well plate and photographed at 7days. For mammosphere assay, 1 × 10^3^ cells/well seeded into 96-well ultra-low adhesion plates in DMEM medium.

### Luciferase reporter assay

3 × 10^4^ cells were transfected BRE-Luciferase reporter plasmid using Lipofectamine 2000 (Invitrogen). The cell lysates collected 48 hr after transfection, and the luciferase activities were measured using the Enhanced Luciferase Assay Kit (BD Biosciences).

### RNA preparation and RT-PCR analysis

Total RNA was isolated using RNeasy Mini Kits (Qiagen), and RT-PCR analysis was performed using a One-Step RT-PCR kit (Qiagen) according to the manufacturer’s instructions.

For quantitative RT-PCR, reverse transcription performed with the Superscript IV First-strand synthesis system (Invitrogen) and amplification performed with SYBR Green Mix I (Roche), and All PCR analyses were conducted in triplicate using the 7900HT Fast Real-Time PCR System (Applied Biosystems). The primer sequences used to amplify the investigated genes listed in Supplementary Table 1.

### Statistical analysis

Data expressed as the means ± SEM. Statistical analyses of the data conducted via the Student’s t-test (two-tailed). Differences were considered statistically significant at *P* < 0.001.

## Acknowledgments

This work supported by a National Research Foundation of Korea grant (2015R1D1A1A01059406, 2018R1C1B6008372 to MSK). Also, This work supported by the Cooperative Research Program for Agriculture Science & Technology Development (Project no. PJ0132772019 to WJ), Rural Development Administration.

## Supplementary Figure Legends

**Supplementary Figure 1.**
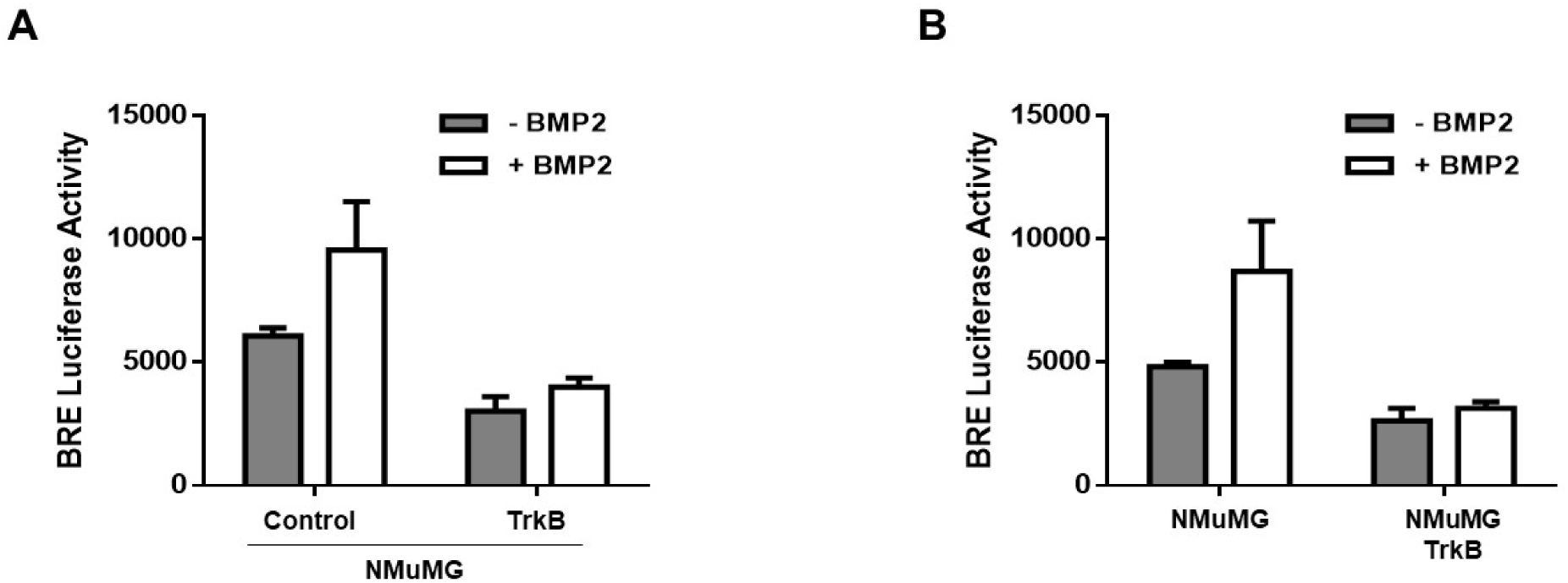
TrkB inhibits BMP signaling. (A) The activity of BMP-2-responsive BRE Luciferase reporter in NMuMG cells transfected with the TrkB. **Control versus treatment with BMP-2, *P* < 0.05. n = 3. (B) The activity of BMP-2-responsive BRE Luciferase reporter in NMuMG and NMuMG-TrkB cells. **Control versus treatment with BMP-2, *P* < 0.05. n = 3.

**Supplementary Figure 2.**
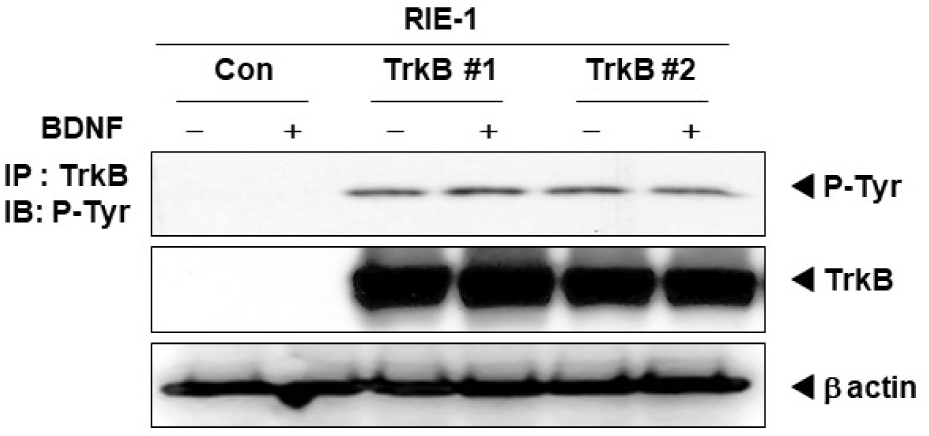
TrkB overexpression leads to the autophosphorylation of TrkB without BDNF. Western blot analysis of the whole-cell lysates and immunoprecipitates of V5 derived from RIE-1 or RIE-1-TrkB cells, as indicated.

**Supplementary Figure 3.**
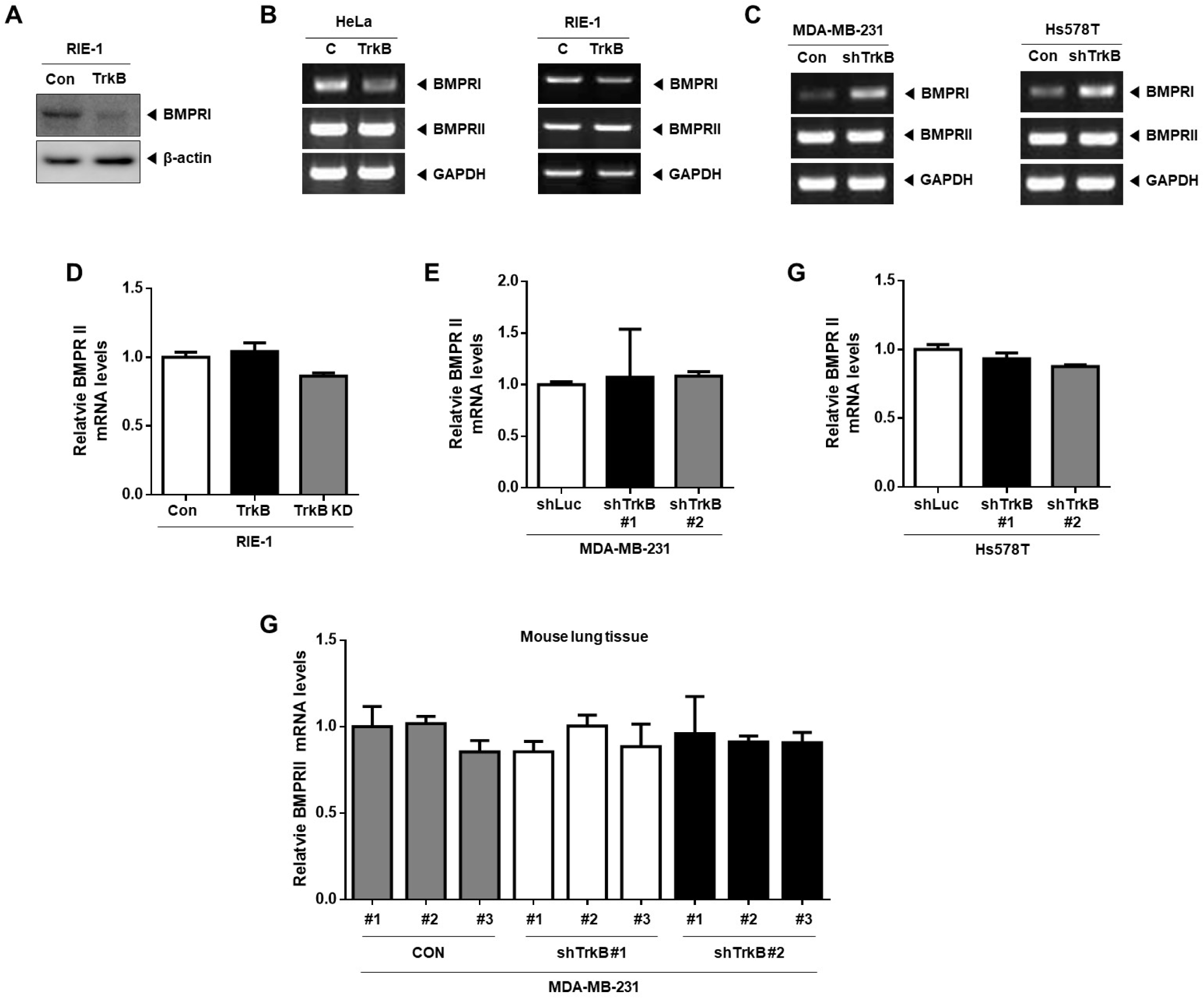
TrkB suppresses BMPRI expression. (A) Western blot analysis of the expression of BMPRI in RIE-1 and RIE-1-TrkB cells. Loading control, β-actin. (B) RT-PCR analyses of BMPRI and BMPRII mRNAs in RIE-1, HeLa, RIE-TrkB, and HeLa-TrkB cells. (C) RT-PCR analyses of BMPRI and BMPRII mRNAs in Hs578T, or MDA-MB-231 control-shRNA and TrkB-shRNA cells. (D) The relative expression levels of mRNA encoding BMPRII in RIE-1, RIE-1-TrkB, and RIE-1-TrkB KD cells, as determined by quantitative RT-PCR. Loading control, 18S. P < 0.05, t-test. (E) The relative expression levels of mRNA encoding BMPRII in MDA-MB-231 control-shRNA and TrkB-shRNA cells, as determined by quantitative RT-PCR. Loading control, 18S. P < 0.05, t-test. (F) The relative expression levels of mRNA encoding BMPRII in Hs578T control-shRNA and TrkB-shRNA cells, as determined by quantitative RT-PCR. Loading control, 18S. P < 0.05, t-test. (G) The relative expression levels of BMPRI mRNA in the lungs of individual mice harboring either MDA-MB-231 control-shRNA or TrkB-shRNA cells, as determined by quantitative RT-PCR. Loading control, 18S. P < 0.05, t-test.

**Supplementary Figure 4.**
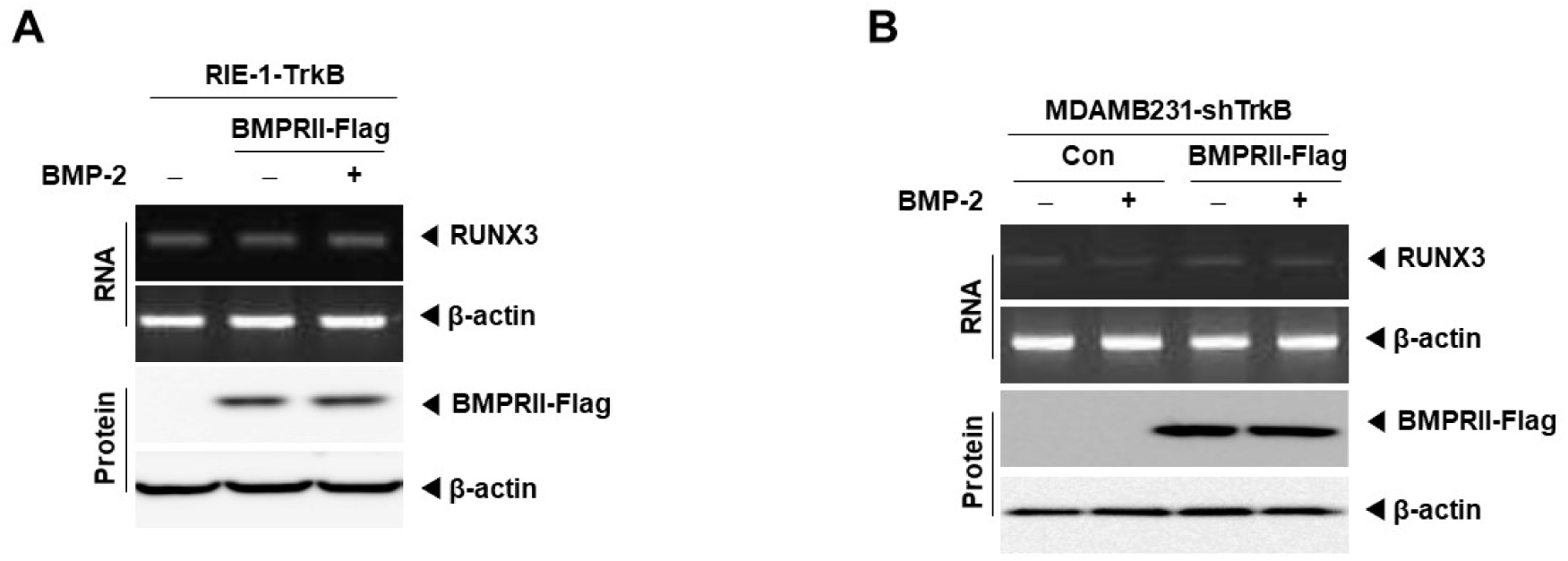
Upregulation BMPRII in the presence of TrkB can not lead to the induction of RUNX3 expression. (A) The relative expression levels of RUNX3 mRNA and protein in RIE-TrkB cells transfected with BMPRII and treated with BMP-2 (I) The relative expression levels of RUNX3 mRNA and protein in MDA-MB-231 TrkB-shRNA cells transfected with BMPRII and treated with BMP-2

